# The effect of parental provisioning on the development of prey preferences in great tit (*Parus major*)

**DOI:** 10.64898/2026.07.03.736371

**Authors:** Linda Nevala, Christopher J Irving, Rose Thorogood, Suvi Ruuskanen, Liisa Hämäläinen

## Abstract

To make adaptive foraging decisions, naive individuals need to gather information about the local prey community. Besides sampling prey personally, the young could gather information about prey profitability by observing the foraging behaviour of other individuals, and parental provisioning provides the first opportunity to acquire this social information. Still, previous research on vertical transmission of prey preferences from parents has provided mixed results that are often confounded with other information sources, such as siblings and peers. It is also not known whether information from parents can change potential innate biases against certain prey types, such as avoidance of warningly coloured insects. Here, we tested whether social information acquired by offspring during parental provisioning influences the development of prey preferences in a generalist predator, the Great Tit (*Parus major*). We brought 15 great tit broods and their parents into captivity at late nestling stage (14 days old) and divided them into three social information treatments where parents were provided with either brown, red or yellow palatable maggots to feed to their dependent young for 8 days. Once foraging independently from parents, we conducted a preference test where juveniles were offered the full array of coloured maggots. Regardless of palatable exposure to typical warning-coloured maggots (i.e. red and yellow), juveniles consistently preferred yellow over red, and preferred brown maggots the most (i.e. lacking warning coloration). This supports the existence of innate biases against typical warning colours, and that social information from parents is unlikely to override these, at least when alternative prey is easily available.

## INTRODUCTION

To make optimal foraging decisions, individual predators are expected to gather information about prey profitability, including nutritional value as well as the costs of ingesting toxic defensive chemicals that the prey might possess (Marples et al., 2018). Innate responses to different prey types (Smith, 1975, 1977; Lindström et al., 1999) can reduce the cognitive load of these foraging decisions, but when environments change (e.g. non-toxic mimics fluctuate in prevalence) it should pay individuals to refine their choices. Learning experiments with independent birds, for example, demonstrate that both personal and social exposure to palatable prey items with typical warning signals can overcome initial biases (Roper, 1990; Teichmann et al., 2020; Hämäläinen et al., 2022). However, it remains unclear whether naive dependent young can take advantage of parents’ knowledge of local prey profitability to update information about prey signals. Previous studies have demonstrated that the diet of juvenile birds differs from more experienced individuals (Coomes et al., 2025), and younger individuals are more likely to make mistakes and attack unprofitable prey (Exnerová et al., 2007, Mappes et al., 2014), suggesting that adaptive foraging decisions require at least some learning. In altricial species, parents provide the first opportunity to observe foraging decisions, and theories predict vertical transmission of information from parents to offspring (McElreath & Strimling, 2008). This social information from parents could potentially shape offspring responses to the local prey community, providing a mechanism for non-genetic inheritance of prey preferences that could influence predator-prey dynamics. Here, we tested this hypothesis by investigating whether prey items provided by parents influence the prey preferences of offspring once they begin foraging independently.

Vertical transmission of information has been demonstrated in many foraging contexts. In some cases, this can be intentional as observed in white-tailed ptarmigans (*Lagopus leucurus*) that call their chicks to nutritionally rich food sources (Allen & Clarke, 2005; Clarke, 2010). More often, parents provide social information about food locations, preferences and foraging techniques unintentionally (Danchin et al., 2004; Dall et al., 2005). Information transfer from parents is particularly common in species that have prolonged parental care (e.g. primates and cetaceans), and it can lead to cultural transmission of feeding preferences and techniques across generations (Whiten, 2021). However, relying on social information provided by parents might not always be adaptive, especially if the environment is changing rapidly (McElreath & Strimling, 2008) or if there is evidence that parents’ foraging behaviour might not be optimal (Boogert et al., 2014). Whether and for how long young individuals use social information from their parents is therefore likely to be context dependent (Kendal et al., 2018). For example, in terms of prey preferences, the value of information from parents could depend on the temporal and spatial stability of the prey community (Seppänen et al., 2007; Hämäläinen et al., 2022).

Social transmission of prey selection has been studied mostly in birds as they are the main predators of many insects where warning coloration is a common adaptation (Ruxton et al., 2018). While there is now good evidence that independently foraging birds can use social information about prey profitability (Mason & Reidinger, 1982; Skelhorn, 2011; Landová et al., 2017; Thorogood et al., 2018; Hämäläinen et al., 2020), the effect of any potential information conveyed by parents during provisioning remains poorly understood (Franks et al., 2019). Cross-fostering experiments with blue tits (*Cyanistes caeruleus*) and great tits (*Parus major*) have shown that cross-fostered individuals shift their foraging niches towards the foster species, particularly in prey size, and that these effects can persist across years (Slagsvold & Wiebe, 2007; 2011). However, whether these feeding preferences were vertically transmitted from foster parents or acquired socially from other individuals remains unresolved (Wild et al., 2025). Another recent study testing “the early learning of the foraging niche hypothesis” did not find a correlation between the type of insects provided to great tits in the nest and the prey that those individuals later provided to their own chicks (Olivé-Muñiz et al., 2025). These contrasting results may reflect the challenge of isolating parental effects, given that great tits are known to flexibly update their foraging behaviour according to changing conditions (Aplin et al., 2017), and juveniles also use social information from their siblings and peers (Wild et al., 2025). In addition, juveniles might use social information only about certain prey traits, with previous studies focusing mainly on prey size (Slagsvold & Wiebe, 2011) or broad classification of insect types (Slagsvold & Wiebe, 2011, Olivé-Muñiz et al., 2025) and less on other traits such as prey colouration (but see a comparison of green and brown larvae in Slagsvold & Wiebe, 2011). To investigate vertical transmission of prey preferences, we therefore need controlled experiments that manipulate prey type and access to information separately from both parents and peers.

Besides acquiring prey preferences socially or through personal sampling, individuals are likely to have innate biases towards and/or against different foods. In the predator-prey context, predators may innately avoid aposematic warning colours, especially when the costs of attacking are high (Smith, 1975, 1977). This has been demonstrated in turquoise-browed motmots (*Eumomota superciliosa*) and great kiskadees (*Pitangus sulphuratus*) that show innate avoidance of typical yellow and red striped colour patterns of coral snakes (Smith, 1975, 1977), and in different species of egrets and herons that innately avoid venomous sea snakes (Caldwell & Rubinoff, 1983). Innate avoidance of typical warning colours such as red and yellow has also been found in domestic chicks (*Gallus gallus domesticus*, Schuler & Hesse, 1985; Roper, 1990; Taylor et al., 2026) and northern bobwhites (*Colinus virginianus*, Mastrota & Mench, 1995). In great tits, hand-reared birds were found to avoid prey with yellow and black stripes (Lindström et al., 1999), but showed no evidence of equivalent innate avoidance of red and black firebugs (Exnerová et al., 2007), indicating that birds might have innate avoidance only against specific colour patterns. Whether social information from parents could change these initial colour biases remains untested, although this could influence predator-prey dynamics. For example, previous studies have demonstrated that social information about defended aposematic prey can reduce predation pressure on warningly coloured prey (Thorogood et al., 2018; Skelhorn, 2011), whereas social information about palatable Batesian mimics can increase the predation pressure on mimics and their defended models (Alcock, 1969; Hämäläinen et al., 2021). Similarly, young individuals might gain social information about warningly coloured but palatable Batesian mimics from their parents and be more likely to attack similar prey themselves, increasing the attack rates on mimics and their models. By uncovering the strength of vertical transmission of prey preferences on foraging decisions, the mechanisms that maintain predator-prey dynamics could be understood.

We tested vertical transmission of prey preferences using great tits as our model predators. Great tits are an ideal study species to answer this question, as they are generalist predators that provide their offspring a wide variety of different insect prey, with the majority of the diet consisting of Lepidopteran larvae (Wilkin et al., 2009), which also include aposematic species (Mappes et al., 2014). We temporarily captured great tit nestlings and their parents when the nestlings were in the late nestling stage, fully dependent on parental feeding (14-15 days old). The parents were provided with either red or yellow maggots to feed to their chicks to investigate if the chicks would develop a preference for the provided colour and be more likely to choose it when foraging independently. Both red and yellow are typically observed in aposematic insects (Ruxton et al., 2018), so we also tested whether social information could change the potential innate wariness of these colours. To test for initial biases against red and yellow without social information, we included an additional treatment where chicks were not exposed to either colour (parents were provided with brown maggots). When the juveniles started sampling food independently (approximately from the age of 25 days onwards), they were not given any maggots, so there was no opportunity to use personal information or social information from their siblings. We then conducted a preference test in which the juveniles were presented with a choice of brown, red and yellow maggots. We predicted that the juveniles would use social information acquired from their parents and therefore show a preference for the same colour that they were fed during the provisioning period.

## METHODS

### Nest monitoring

The experiment was conducted during the spring and summer of 2025. We had approximately 400 nestboxes located around Jyväskylä, Central Finland (62°N 25°E), which were monitored once a week throughout the nest building and incubation stages in April and May. We had 65 great tit pairs nesting in the boxes, but some nests failed (7 nests) or were not used in the experiment for logistical reasons (19 nests), so for this experiment, we used chicks from 39 nests. The chicks were ringed and measured at 7 days old, and we collected a small blood sample (10μl) for molecular sexing that was done following previously established protocols (Aljanabi & Martinez, 1997; Griffiths et al., 1998).

The experiment was conducted simultaneously with another project that aimed to investigate the effect of early-life gut microbiome on great tit physiology and behaviour (Irving et al., in prep.). For this reason, the nests were divided into three gut microbiome manipulation treatments where the chicks were fed with a daily dose of i) 100mg/kg antibiotic (amoxicillin, Amovet vet), ii) prey chemical defences (two pyrrolizidine alkaloids: 6 μg Senecionine and 60 μg Seneciphylline), or iii) water (120μl, control). The treatment was given to the nestlings four times at ages 7, 9, 11 and 13 days by pipetting the solution into their beaks. This treatment was not related to our study question on social information use, and it is described in more detail elsewhere (Irving et al. in prep.). However, we considered the microbiome treatment in our study design when we divided the nests into social information treatments (see below). We also included the microbiome treatment in the statistical analyses to test for any potential effects of the microbiome manipulation on birds’ colour preferences. While this additional treatment introduced another covariate in our study, it also allowed us to conduct multiple experiments with the same individuals which is an important ethical consideration given the requirements of captivity.

### Raising birds in captivity

When the nestlings were at late nestling stage (14–15 days old) 90 nestlings and 30 parents were captured and transferred to Konnevesi research station, where they were raised in captivity. This was done by forming 15 foster nests that each had six nestlings (Figure 1). Each foster nest was formed by combining nestlings from 2–4 different nests (1–4 nestlings/nest) where the nestlings were of the same age (±1 day) and from the same microbiome treatment. We aimed to get as balanced a ratio of both sexes as possible (48 females, 40 males, 2 unknown). For each foster nest, we captured one pair of parents, and the nestlings that were not taken into captivity were divided among the remaining nests. Because we took the birds into captivity when the nestlings were two weeks old, we could not control the parental provisioning and social information that the nestlings received before this. However, all nestboxes were in similar habitats, which should reduce variation in prey items provided by the parents, and this variation should be similar in all social information treatments. Furthermore, great tit nestlings open their eyes when they are 8–9 days old, so the chicks were unable to gather any visual cues about prey before that.

**Figure 1.**
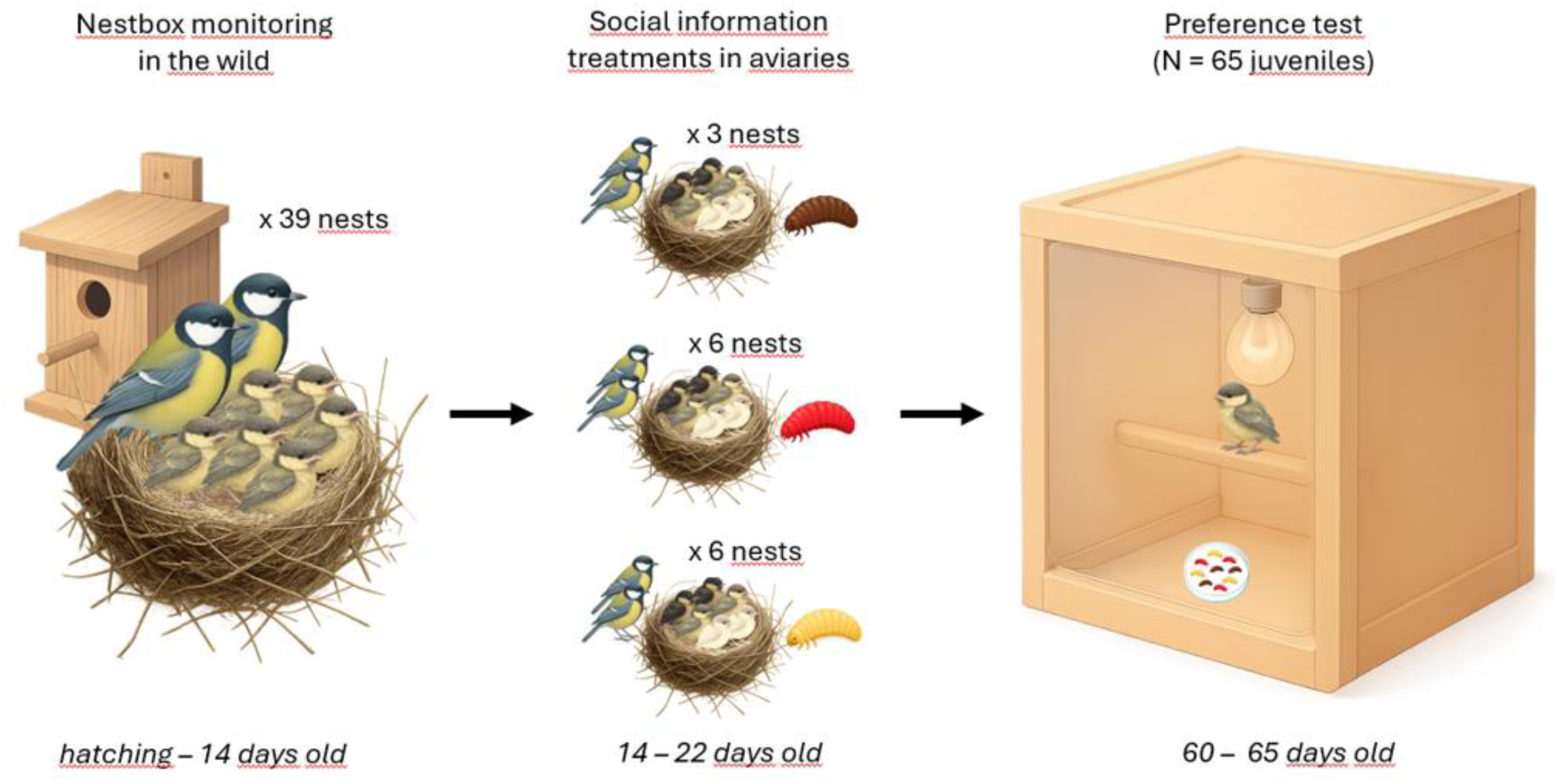
Experimental design. The nestboxes were monitored in the field until the chicks were 14 days old. At the age of 14–15 days, nestlings from 2–4 nests (illustrated here by differently coloured chicks) were combined to form 15 foster nests that each had six nestlings. The foster nests and their parents were brought into captivity (separate aviaries), where the parents were provided with either brown (3 broods), red (6 broods) or yellow (6 broods) maggots to feed to their chicks for eight days. When the juveniles were 60–65 days old, they underwent a preference test where they were presented simultaneously with three brown, three red and three yellow maggots and the order of their choices was recorded.

In captivity, each foster brood and its parents were housed in a large outdoor aviary (2 × 2 × 2 m) that had double wire mesh (1 × 1cm) walls. The cages included many fresh tree branches that provided cover and perching places, and the floor was covered with wood chips. Fresh water and food were always available. The birds were provided with a diverse diet of live insects (mealworms, giant mealworms, wax moth larvae, soldier fly larvae, crickets) and dried insects (dried mealworms, crickets and silkworms), and sunflower seeds, peanuts and tallow. Fresh vegetables were provided occasionally as enrichment. Although the cages were next to each other, there were four layers of wire mesh walls between them and many tree branches on the sides of the cages, so it is unlikely that the chicks were able to observe the prey choices of the birds in the neighbouring cages.

The chicks fledged approximately at 18–20 days old. After that, parents continued feeding them for 2–3 weeks. Parents were released back at their capture site when the chicks were 50 days old, which ensured that all chicks were completely independent and foraging efficiently. The juveniles were kept in captivity for three months to conduct behavioural assays. Unfortunately, 27 juveniles died during this captivity period, most likely due to various infections (see Ethical note). The rest of the juveniles (*N* = 63) were released to the Konnevesi research station surroundings in August. They were provided supplementary food near the cages for the first week after the release and feeding continued at the research station throughout the autumn and winter.

### Social information treatments

When the broods were brought into captivity, they were divided into three social information treatments where the parents were provided with 1) brown maggots (3 broods, 18 nestlings), 2) red maggots (6 broods, 36 nestlings) or 3) yellow maggots (6 broods, 36 nestlings, Figure 1). The microbiome manipulations were balanced across these treatments. Our main study question was to investigate whether chicks use social information about yellow or red maggots from their parents, and brown maggots were included to test for potential biases towards these two warning colours when the chicks did not receive social information about either colour. The number of nests that received social information about brown maggots was therefore lower than in the other two treatments.

Red and yellow fly maggots were commercially available, and brown maggots were prepared by colouring white maggots (all ordered from Onkitukku, JIG, Finland) with brown food dye (Colour Mill Aqua Blend, chocolate). The food dye was odourless and flavourless, but to ensure that it would not influence birds’ preferences, we also used the respective food dye to cover the yellow (Colour Mill Aqua Blend, yellow) and red (Colour Mill Aqua Blend, red) maggots. Each nest was provided with 10g of coloured maggots per day in a white bowl, and the colouring was done just before presenting the maggots to the birds. While we could not confirm that the parents were feeding an equal number of maggots to each chick, we observed the parents feeding the maggots to the chicks in all colour treatments, and all provided maggots were consumed during the same day they were offered.

We began providing coloured maggots on the same day birds were brought into captivity (when the chicks were 14–15 days old) and continued until chicks were 22 days old (i.e. soon after fledging). This relatively short period of provisioning was because the chicks started to sample food independently when they were approximately 25 days old, and we needed to avoid chicks having opportunities to gain information about the coloured maggots independent from parental provisioning. Because we did not know the exact age when the chicks would start foraging independently, the first two nests received maggots until the chicks were 26–30 days old. It is therefore possible that these chicks sampled some maggots themselves. To be conservative, we repeated analyses excluding these individuals, but this did not change our conclusions (see Supplementary material).

### Preference test

When the chicks were 60–65 days old (mean = 62 days), we conducted a preference test to see if their foraging choices were influenced by social information from the parents. At this age, the chicks had been foraging independently for approximately one month without access to any maggots, so we tested their short-term prey preferences. We tested a total of 80 chicks, but 15 chicks did not eat the maggots in the test conditions and were not used in the analysis. Our final sample size was therefore 65 chicks that had received from their parents 1) brown maggots (*N* = 12: 2 males, 9 females, 1 unknown), 2) red maggots (*N* = 26: 11 females, 15 males), or 3) yellow maggots (*N* = 27: 12 females, 15 males). There were no differences between sexes or among social information or microbiome treatments in birds’ likelihood to complete the test (see Supplementary material).

The birds were moved to individual test cages (50 × 66 × 49 cm) in the morning of the testing. The test cages were made of plywood and had a plexiglass front wall that allowed us to observe the birds. The cages were illuminated with a light bulb (LED 4.5 W, 4000 K, 360°, 470 Lumen, Airam Otso) and included a perch. Before the test, the birds were allowed to habituate to the cages for at least 90 minutes. Water was always available, but food was restricted before and during the test to motivate the birds to forage.

In the test, the birds were presented with three brown, three red and three yellow maggots (Figure 1). The maggots were randomly distributed on a petri dish (9 cm diameter) with a white background (white paper glued on the bottom of the dish). To prevent the maggots from moving from the dish, they were euthanised just before offering them to the birds. We recorded the order in which the birds chose to consume the maggots and filmed the assays with a camcorder (Sony Handycam HDR-CX405) to verify the foraging choices later if necessary. To score as “consuming”, the birds were required to eat the entire maggot. If the bird did not eat any maggots within 30 minutes, or there had been 15 minutes since eating a maggot, the petri dish was removed, and the bird was given a break (at least 30 minutes) before continuing the test with the same petri dish. When necessary, birds were given mealworms during these breaks to ensure that the food restriction time was not longer than 90 min. The test was completed when the bird had eaten all nine maggots. Most birds (*N* = 59) completed the test in one day. If the bird did not eat the maggots during five 30-minute attempts, the test was finished (*N* = 21). When logistically possible, the birds that did not complete the test during the first day were tested again 2–3 days later (*N* = 9). The test was restarted with all nine maggots, as only two of the retested birds had eaten maggots during the first day. One of these birds ate one brown maggot on the first day but chose a yellow maggot first when retested. In this case, we considered brown as a first choice, but we calculated the preference scores (see below) based on the second test day. Of the nine birds, six finished the test on the second test day.

### Ethical note

The study was conducted with permits from the Finnish Project Authorisation Board (ESAVI/8016/2025) and Central Finland Centre for Economic Development, Transport and Environment (VARELY/1344/2025). The parent birds were caught using traps at the entrance of the nestbox. The trapping was continued for a maximum of 90 minutes, with checks every 15 minutes, and we did not observe any adverse effects from catching on the parents or nestling survival. The birds were housed in family groups (parents and six chicks) in outdoor aviaries which were cleaned regularly. The aviaries included many fresh tree branches that were replaced when necessary, and the birds were provided with enrichment boxes that included mealworms and crickets to encourage natural food searching behaviour. All parents (*N* = 30) were released back at their capture site in good condition after one month of captivity. During the behavioural assays with juveniles, short food restriction was necessary to motivate the birds to forage, but this was kept as short as possible (maximum of 90 minutes) and the birds were provided with preferred food (mealworms) after the tests. While the behavioural tests might have caused short-term stress to the birds, there was no evidence that they affected the birds’ body condition or survival. The juveniles were kept in captivity for three months to conduct the preference tests, as well as other behavioural assays related to a simultaneous experiment (Irving et al., in prep.). Unfortunately, 27 juveniles died during this captivity period, most likely due to infections that occurred during the moulting period and were possibly exacerbated by a two-week heatwave in July. Laboratory analyses identified coccidiosis as a likely cause of these deaths, and after consulting the veterinarian, birds that displayed any symptoms (inactive, weight loss) were treated with medicine (Baycoxine). The rest of the juveniles (*N* = 63) were released in August. The survival of great tit fledglings in the wild is typically very low, with approximately 50% mortality during the first weeks after fledging (Naef-Daenzer et al., 2001), so despite the unfortunate mortality in our experiment, the overall survival of juveniles was higher than expected in natural conditions.

### Statistical analyses

We first analysed whether birds’ first prey choice (brown, red or yellow maggot) was influenced by their social information treatment (parents providing brown, red or yellow maggots) using a G-test. We also used G tests to investigate if other covariates, birds’ sex and the gut microbiome treatment, influenced their first prey choices. To analyse whether birds preferred one colour regardless of the social information treatment, we used a binomial test to compare the birds’ first choices to the predicted equal probability of choosing any of the three colours.

To analyse preferences across the whole foraging assay, we calculated a preference score for each colour (Taplin, 2007). This was done by first ranking the maggots from 1 to 9, based on the order in which birds ate them. We then calculated an average rank for each of the three colours for each bird, with smaller values indicating that birds preferred that colour. This preference score generated values ranging from 2 (all maggots of one colour were consumed before any other colours, i.e. (1+2+3)/3) to 8 (all maggots of one colour were consumed last, i.e. (7+8+9)/3), allowing us to compare relative preferences for each colour. This was done using linear mixed effects models, where the preference score was included as a response variable that could be modelled using a Gaussian error distribution. To investigate whether colour preference was influenced by social information from parents, the fixed effects included an interaction term between social information treatment (parents providing brown, red or yellow maggots) and maggot colour (brown, red or yellow). We also ran separate models that included an interaction term between sex and maggot colour to test whether this affected preferences, and between microbiome treatment and maggot colour to check for any unexpected effects of the simultaneous experiment. Diagnostic tests and residual plots indicated that the linear model was appropriate. Because of heteroscedasticity among maggot colour groups, we allowed different variances for each group using the *nlme* package (Pinheiro & Bates, 2025). Models were initially built with a hierarchical random-effects structure, where bird identity was nested within the original nestbox (in the field) and the cage (i.e. foster brood) in captivity. However, nestbox and cage were excluded from the final model because their variances were very small (< 0.0001) and removing them improved model fit based on Akaike’s information criterion. Bird identity was left in the model to control for repeatable measures from each bird (i.e. preference score for each colour). Simplifying the random effect structure to include only bird identity did not change the estimates of the fixed effects. The final terms in the model were selected based on their significance and AIC values.

All analyses were conducted in R version 4.4.3 (R Core Team 2025). The significance of the main effects in the models was tested using the Anova() function in the *car* package (Fox & Weisberg, 2019), and pairwise post-hoc comparisons were conducted using the *emmeans* package (Lenth, 2025). The model assumptions were checked visually and by using the *DHARMa* package (Hartig, 2024), and the graphs were generated using the *ggplot2* package (Wickham, 2016).

## RESULTS

### First choice

The birds consumed a brown maggot as their first choice more often than predicted by chance (binomial test: 52/65, *P* < 0.001). This preference did not differ among social information treatments (G-test: *G* = 5.662, *df* = 4, *P* = 0.23; Figure 2), or between males and females (G-test: *G* = 1.406, *df* = 2, *P* = 0.50). There was also no effect of the early-life gut microbiome treatment on birds’ first choices (G-test: *G* = 3.760, *df* = 4, *P* = 0.44).

**Figure 2.**
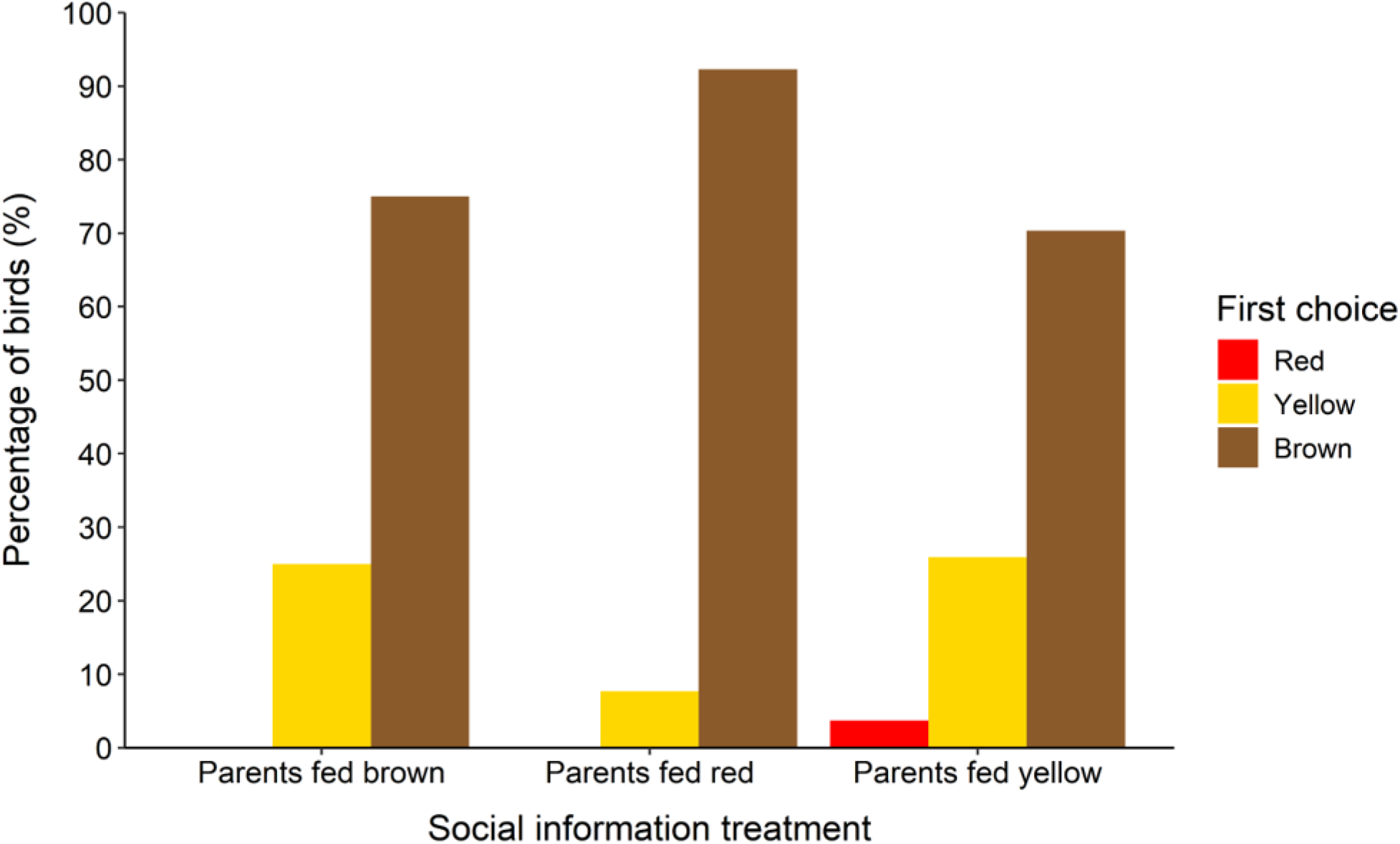
Birds’ (*N* = 65) first foraging choices in the preference test. The bars show the percentage of the birds choosing a red, yellow or brown maggot as their first choice in each social information treatment (parents providing brown maggots: *N* = 12, parents providing red maggots: *N* = 26, parents providing yellow maggots: *N* = 27).

### Overall preference

Social information from parents did not influence birds’ preference for coloured maggots during the foraging test (Table 1). Regardless of the social information treatment, there was a significant effect of maggot colour on the foraging choices (Table 1), with birds preferring brown maggots over red (brown vs. red: estimate = –4.150 ± 0.234, *t* = –17.787, *P* < 0.001) and yellow (brown vs. yellow: estimate = –2.460 ± 0.279, *t* = –8.828, *P* < 0.001; Figure 3). Birds also had a significant preference for yellow over red maggots (yellow vs. red: estimate = –1.690 ± 0.273, *t* = –6.195, *P* < 0.001; Figure 3). Neither birds’ sex nor early-life microbiome treatment influenced their colour preferences (Table 1).

**Figure 3.**
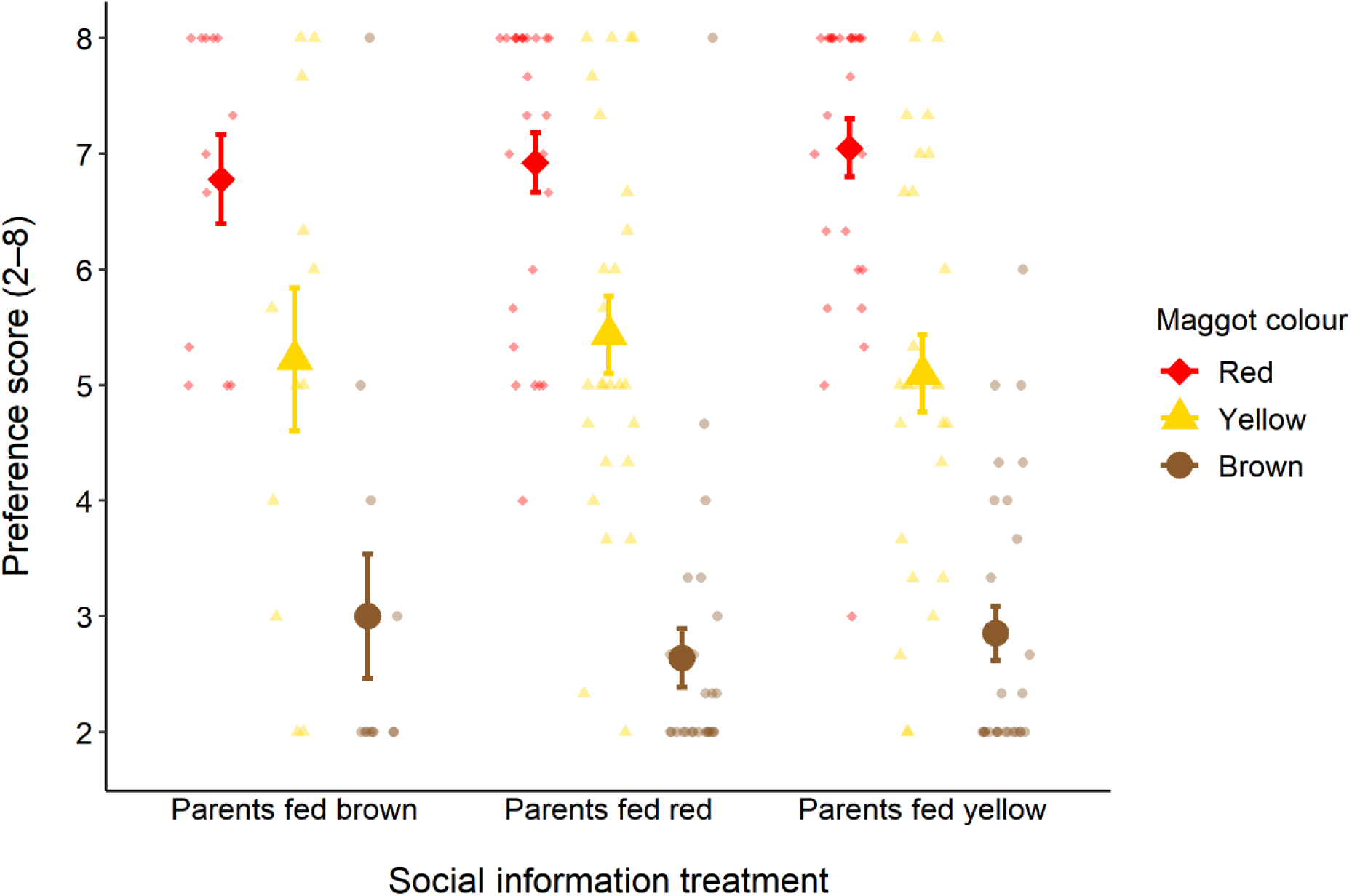
Birds’ (*N* = 65) preference scores for red, yellow and brown maggots in the preference test. Preference scores indicate the order in which the birds consumed the maggots, with smaller scores meaning that birds preferred that colour. Big symbols present the mean (± s.e) preference scores for the three colours in each social information treatment (parents providing brown maggots: *N* = 12, parents providing red maggots: *N* = 26, parents providing yellow maggots: *N* = 27). Small symbols show individual variation.

**Table 1.** Comparison of linear mixed effects models explaining the preference scores. The table shows the main effect test statistics for maggot colour (final model on top) or for interactions between maggot colour and other variables, and AIC values for each model. All models included bird identity as a random effect.

| Terms in the model | $\chi^2$ | *df* | *P* | Model AIC |
| --- | --- | --- | --- | --- |
| ~ colour (final model) | 317.51 | 2 | < 0.001 | 718.48 |
| ~ colour × sex | 2.148 | 4 | 0.71 | 720.81 |
| ~ colour × social information treatment | 1.412 | 4 | 0.84 | 727.89 |
| ~ colour × microbiome treatment | 0.917 | 4 | 0.92 | 728.90 |

## DISCUSSION

Inexperienced juvenile predators need to gather information about the profitability of different prey types, and this can be acquired by sampling prey directly (Skelhorn et al., 2016) or by learning from the foraging choices of others (i.e. social information use; Hämäläinen et al., 2022). Here, we tested whether prey items provided by parents influence prey preferences of juvenile great tits once independent. Contrary to our prediction that the juveniles would prefer the familiar colour provided by parents, we found no evidence that prey colour preferences were transmitted vertically. Instead, the juveniles preferred brown prey overall, and yellow over red, regardless of the social information they received. These results support the idea that avian predators have innate avoidance against typical warning colours (Smith, 1975, 1977; Schuler & Hesse, 1985; Lindström et al., 1999) and suggest that these innate biases may have a stronger influence on early-life foraging decisions than social information acquired during the parental provisioning period.

While previous studies with great tits have investigated whether vertically transmitted prey preferences last across years (Slagsvold & Wiebe, 2007, 2011; Olivé-Muñiz et al., 2025), we tested the effect of parental provisioning on the prey choices of two-month-old juveniles. The value of social information is predicted to decrease with time as ecological conditions change (Seppänen et al., 2007). For example, prey community composition is likely to vary over time and space, so social information received from parents might quickly become outdated. We therefore predicted vertical transmission of social information to be stronger early in life, compared to longer-lasting preferences across years, but we found no evidence to support this prediction. Predators may value information about some prey traits more than others. Because colouration often provides information about prey profitability (Aronsson & Gamberale-Stille, 2008; Ruxton et al., 2018), we predicted that juveniles would copy colour preferences from their parents; however, it is possible that other traits, such as prey size, type, or specific signal pattern, provide more valuable information about prey quality. Social information use often varies among individuals (Mesoudi et al., 2016), and the observed variation in preference scores, especially in the treatment where the chicks received yellow maggots from their parents (Figure 3), suggests that some individuals might have copied their parents even though we did not find overall differences among treatments. However, we did not see any clear patterns that these individual differences would have been explained by rearing conditions (i.e. the original nestbox in the field or the foster brood in captivity) or the sex of the bird.

Previous studies with great tits (Wild et al., 2025) and other bird species (Franks et al., 2020) have shown that juveniles often switch to social information from their siblings or peers instead of parents. Opposite to previous field studies that investigated the correlation between the diet received by great tit nestlings and the one they provided to their own chicks (Slagsvold & Wiebe, 2007, 2011; Olivé-Muñiz et al., 2025), we conducted our experiment in controlled captive conditions where we could prevent social information use from peers by offering coloured maggots only when the chicks were not yet foraging independently. However, even when there was no other information source about the maggot colour, the birds still ignored social information from their parents. This could be explained by relatively short exposure to the maggots, as we only provided them when the chicks were 14–22 days old. It is not known if a certain time period during great tit development is particularly important for acquiring social information. The nestlings open their eyes when they are 8–9 days old, but the colour of the prey might be hard to see in the dim light conditions inside the nestbox, so the prey items provided after fledging could provide better opportunities to gather social information. Chicks might be most likely to copy their parents’ prey choices when they start sampling prey independently but are still partly fed by their parents; however, disentangling personal and social information during this time period is challenging. One limitation of our study is that we could not record how frequently the parents fed the maggots to their chicks and whether the maggots were equally divided among the chicks in the same aviary. Nevertheless, we confirmed that the parents were feeding some of the maggots to the chicks in all social information treatments, and all maggots were consumed. This means that even if parents did not provide coloured maggots to each chick, all fledged chicks should have had opportunities to observe their parents feeding other chicks or eating the coloured maggots themselves.

Regardless of the provided social information, all chicks preferred brown, whereas red was consistently consumed last. This supports the idea that avian predators can have innate avoidance of yellow and red that are typical warning colours of aposematic prey (Schuler & Hesse, 1985; Roper, 1990; Mastrota & Mench, 1995; Lindström et al., 1999). In contrast, a study with hand-raised great tits did not find innate avoidance of red and black aposematic firebugs (Exnerová et al., 2007), and in the field, predation risk for warningly coloured larvae is highest when inexperienced fledglings are abundant, suggesting that avoidance of specific warning signals needs to be learned (Mappes et al., 2014). The response to the colours is also likely to be context-specific, with a previous study finding six-month-old great tit juveniles preferring red almond flakes over green, while this preference was not observed in older birds (Teichmann et al., 2020). This supports the idea that colour preferences depend on the food type (i.e. insect vs. fruit, Gamberela-Still & Tullberg 2001, 2007) and are shaped by an individual’s previous experience (Teichmann et al., 2020). Interestingly, we found a clear preference for yellow over red maggots, although both are considered typical warning colours (Ruxton et al., 2018). Similar preference for red over yellow was observed in northern bobwhites (Mastrota & Mensch, 1995), and red may be innately perceived as a stronger warning signal than yellow. Importantly, our experiment tested the relative preference of the three colours, so the birds did not completely avoid red prey. All prey were also presented simultaneously, and birds might have attacked red prey more quickly if they did not have alternative prey, as in Exnerová et al. (2007), where different prey types were presented sequentially. Finally, in our experiment, all prey were equally detectable from the white background, and there was no competition from other predators, whereas in nature search costs of cryptic prey and competition with other predators limit prey availability.

When considering innate avoidance, one limitation of our study is that we do not have information about the prey items provided by the parents before day 14 when we took the broods into captivity. Even though the chicks were able to gather visual cues only after opening their eyes on day 8–9, preference for brown maggots may be partly explained by parents providing the chicks with similar-looking brown larvae. Only approximately 15% of Finnish Lepidopteran larvae are estimated to have features of warning colours, with less than 5% having large conspicuous signals (Mappes et al., 2014), so the majority of the diet provided to the nestlings in the wild was likely to consist of cryptically coloured species such as brown larvae. In addition, the diet in captivity included brown mealworms, and even though these were a lighter shade of brown and a different shape than the maggots, they might have influenced chicks’ colour preferences. Exposure to differently coloured prey before day 14 is less likely to explain our finding of juveniles preferring yellow over red because these colours are less common in insects in the area (Chinery, 2012). Although there are several species that include some yellow or red patterns (e.g. many Heteroptera and Coleoptera), it is not known if and how often great tits provide these to their offspring, and how information about colouration is generalised to different patterns and prey types. Finally, while it is unlikely that the parents provided the nestlings with many chemically defended prey, it is possible that the chicks received some negative social information about red and/or yellow prey being unpalatable, which could have later influenced their preferences. Further studies that control the diet during the whole parental provisioning time are therefore needed to confirm that the observed avoidance of red and yellow is innate.

In conclusion, we did not find evidence that social information from parents influenced the foraging decisions of independently foraging great tit juveniles. This might be explained by a conflict between the provided social information and birds’ innate avoidance of warningly coloured insects, and future studies could explore this by conducting a similar experiment with more neutral prey, such as prey with artificial symbols that are evolutionary novel to the birds (Alatalo & Mappes, 1996). It is also possible that the birds ignored social information in our experiment because alternative prey was easily available and information from the parents could be more valuable for improving foraging success in the wild. Juveniles might, for example, form search images of the same prey type provided by their parents, which could increase detection rates of that prey, but this idea remains untested. Previous studies with great tits suggest that preference for prey size might be acquired socially from the parents, but evidence of vertical transmission of prey types is less clear (Slagsvold & Wiebe 2011), possibly because accurate classification of the diet in the wild is challenging. Recent technical developments with metabarcoding that allow identification of the bird’s diet to the insect species level (e.g. Rytkönen et al., 2019; Coomes et al., 2025) could provide a useful tool to quantify whether the diet provided as a chick correlates with the foraging choices of the adult bird. If so, this could have implications for predator-prey dynamics. For example, social transmission of prey preferences could increase predation pressure for a specific prey phenotype and lead to selection favouring polymorphism in prey appearance (Hämäläinen et al., 2022). Investigating how foraging decisions are shaped by a predator’s innate biases and personal experiences, as well as social information acquired from parents and peers, can therefore help us understand predator-prey coevolution and the complex selection pressures for prey defences.

## ACKNOWLEDGEMENTS

We thank Oihana Garcia Carrillo, Maria Correia, Charli Davies, Juho Hartikka, Stijn Kouwenberg, Ella Ruutu and Janna Savin for the help with fieldwork, and Apolline Maitre for sexing the chicks. We also thank Toni Laaksonen for feedback on the writing and staff at the Konnevesi research station for providing facilities for the experiment. The project was funded by the Academy of Finland Research Fellowship (#355869) and a Horizon Europe Marie Sklodowska Curie Actions Fellowship (#101151042) awarded to LH and European Research Council Consolidator grant (#101124827) to SR.

