## Supplementary material for "The effect of parental provisioning on the development of prey preferences in great tit (*Parus major*)"

### Birds' likelihood to complete the preference test

We tested colour preferences of 80 juvenile birds. 59 birds completed the test during the first day (i.e. consumed all 9 maggots). 21 birds did not complete the test during the first day, with 17 of them not eating any maggots, two birds eating one maggot, and one bird eating three and one six maggots. One bird consumed all brown and yellow maggots but did not eat the red ones, and we considered that it had completed the test, as we could calculate the preference score for each colour (i.e. red getting the highest score of 8, indicating that red maggots were least preferred). Similarly, one bird left the last yellow maggot uneaten, and we calculated the preference score for yellow so that we considered it to be chosen last. Whether or not these birds were included in the analysis did not change the results.

We used G-tests to investigate if a bird's sex, social information treatment or gut microbiome treatment influenced their likelihood to complete the test during the first day. There was no difference between females (29/43 completed) and males (29/36 completed) in the likelihood to complete the test during the first day (G-test: $G = 1.758$, $df = 1$, $P = 0.18$). Similarly, social information treatment did not influence birds' likelihood to complete the test (G-test: $G = 4.027$, $df = 2$, $P = 0.13$), with 10/18 birds that were provided with brown maggots, 22/29 birds that were provided with red maggots and 27/33 birds that were provided with yellow maggots completing the test during the first day. Finally, there was no effect of microbiome treatment on birds' likelihood to complete the test (G-test: $G = 1.952$, $df = 2$, $P = 0.38$), with 23/29 birds from the antibiotic treatment, 18/28 birds from the control treatment and 18/23 birds from the prey chemical defence treatment completing the test during the first day. Nine of the 21 birds that did not complete the test during the first day were tested again 2–3 days later, and six of these birds completed the test during the second attempt. All birds were not re-tested because of logistical reasons.

**Supplementary results: excluding the first two nests**

The first two foster nests received maggots for longer than the others (until the chicks were 26–30 days old) because we were not sure when the chicks would start to forage independently. After observing the chicks' behaviour in the first two nests, we confirmed that the chicks started to sample food when they were approximately 25 days old, and we decided to offer coloured maggots for the rest of the nests until the chicks were 22 days old. To test whether longer exposure to the maggots would influence the results, we conducted the analysis excluding the birds ($N = 8$) from the first two nests. This did not change our conclusions, with no significant effect of social information treatment on birds' colour preference scores ($\chi^2 = 0.573$, $df = 4$, $P = 0.97$), and birds preferring brown over red (estimate = –4.120 ± 0.254, $t = –16.236$, $P < 0.001$) and yellow (estimate = –2.260 ± 0.302, $t = –7.498$, $P < 0.001$), and yellow over red (estimate = –1.860 ± 0.288, $t = –6.452$, $P < 0.001$).
